# Sperm mosaicism predicts transmission of *de novo* mutations to human blastocysts

**DOI:** 10.1101/2022.03.28.486034

**Authors:** Martin W. Breuss, Xiaoxu Yang, Valentina Stanley, Jennifer McEvoy-Venneri, Xin Xu, Arlene J. Morales, Joseph G. Gleeson

**Affiliations:** Rady Children’s Institute for Genomic Medicine, San Diego, CA 92123, USA; Department of Neurosciences, University of California, San Diego, La Jolla, CA 92093, USA; Department of Pediatrics, Section of Clinical Genetics and Metabolism, University of Colorado School of Medicine, Aurora, CO 80045, USA; Fertility Specialists Medical Group, San Diego, CA 92123, USA

## Abstract

*De novo* mutations underlie individually rare but collectively common pediatric congenital disorders. Some of these mutations can also be detected in tissues and from cells in a parent, where their abundance and tissue distribution can be measured. We previously reported that a subset of these mutations is detectable in sperm from the father, predicted to impact the health of offspring. Here, in three independent couples undergoing *in vitro* fertilization, we first assessed male gonadal mosaicism, then assessed the transmission of the mutations to their preimplantation blastocysts. We found an overall predictable transmission but slight under-transmission of mutations to blastocysts based upon measured mutational abundance in sperm, and we replicated this conclusion in an independent family-based cohort. Therefore, unbiased preimplantation genetic testing for gonadal mosaicism may represent a feasible approach to reduce the transmission of potentially harmful *de novo* mutations, which could help to reduce their impact on miscarriage and pediatric disease.

## Introduction

Genomic mosaic mutations—present in some but not all cells within a tissue—record the history of embryonic development, environmental exposure, and have a wide range of implications for human health^1,2^. Mosaic mutations are commonly recognized in cancers or localized overgrowth disorders, such as Proteus, CLOVES, and hemimegalencephaly syndromes^3-5^. Increasingly recognized in more complex diseases such as autism spectrum disorder or structural abnormalities^6,7^, mosaic mutations are typically restricted to the individual in which they arise unless they appear prior to the embryonic separation of somatic and germline lineages or within germ cell progenitors. In these cases, gonadal mosaic mutations (comprising gonad-specific and gonosomal) have the potential to transmit to offspring. These will appear as a considerable portion of *de novo* mutations and may result in miscarriage or a congenital or complex disease—often without phenotypes in the parents^8,9^.

We and others previously demonstrated that 5-20% of identified pathogenic *de novo* mutations in a child are detectable in parental tissues, with ejaculated sperm demonstrating the highest rate of occurrence^9-11^. Every male harbors up to dozens of such mutations in sperm, which—in contrast to other paternal mutation types that increase with age^12^—contribute a life-long threat of transmission largely independent of paternal age^13^. As such, they are thought to explain, in part, the individually rare but collectively common risk of congenital disorders from *de novo* mutations^14^. Yet, experimental evidence of transmission to a conceptus of *in situ* identified gonadal mosaic mutations—in contrast to the detection of gonadal mosaicism of already transmitted variants—is lacking. This is a critical point upon which clinical implementation hinges because procedures like preimplantation genetic testing (PGT) could be used to select embryos absent for potentially damaging mutations detected in sperm. Here, we demonstrate that gonadal mosaic mutations detected in sperm from individual males can transmit to their preimplantation blastocysts. We decided to assess this early time point, as it avoids any potential bias introduced by a possible selection of mosaic mutations or their lineages during implantation or survival.

## Results and Discussion

We recruited three couples undergoing *in vitro* fertilization (IVF) for infertility (F01, F02, and F03), where at least eight blastocysts each were donated for research (Figure 1 - table supplement 1). DNA was extracted from each sample, including paternal sperm, using standard procedures^10^, and blastocysts underwent whole genome amplification. Tissue-specific and -shared mosaicism was determined for each donor (F01-F03) in sperm and one somatic tissue (i.e., blood or saliva) as described^13^ (Figure 1a). Whole-genome sequencing (WGS) of sperm samples to 300× read depth allowed ‘best practice’ detection of mosaicism^13^. The computational pipeline demonstrates 90% specificity and >95% sensitivity for mosaic mutation detection at allelic fractions (AF) above 0.03 (Ref ^13^). All putative mosaic mutations and additional control single-nucleotide polymorphisms (SNPs) from the sperm donors were then subjected to validation using massive parallel amplicon sequencing (MPAS, AmpliSeq for Illumina Custom DNA Panel)^15^. This was done for both tissues of the sperm donor, one somatic tissue of the egg donor if provided (F01 and F02), and all available blastocysts.

**Figure 1.**
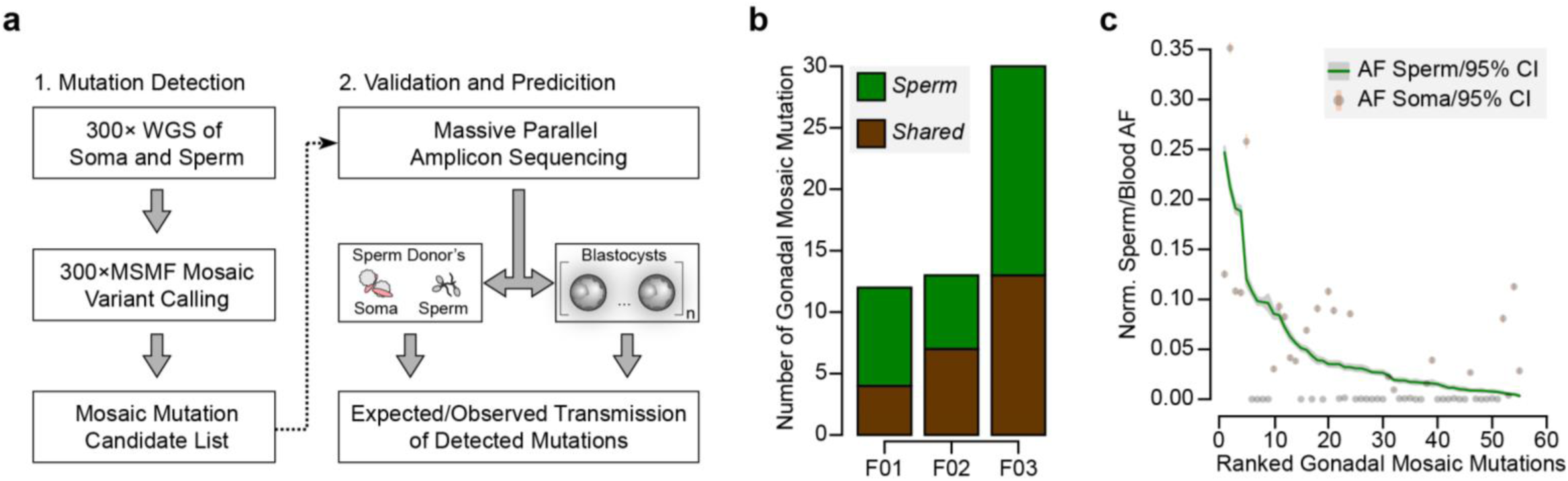
Detection of gonadal mosaicism in three sperm donors. **(a)** Overview of the employed workflow from mutation detection to validation and prediction. A single massive parallel amplicon sequencing (MPAS) panel was used for both detections of mutations in parental tissues and in preimplantation blastocysts. WGS: whole genome sequencing; 300×MSMF: variant calling pipeline on 300× WGS data using MuTect2, Strelka2, and MosaicForecast^13^. (b) Number of MPAS-validated gonadal mosaic mutations for each sperm donor, distinguished by color into sperm-specific (‘*Sperm’*, green) and tissue-shared (‘*Shared’*, brown) mutations. (c) Ranked plot of all gonadal mosaic mutations across the three sperm donors. For each variant, both the allelic fraction (AF; normalized to chromosome count) of the mutation in sperm (green line) and in the soma (brown dot) are shown together with their 95% exact confidence interval. *Shared* mutations tend to be of higher AF compared to *Sperm*.

The three sperm donors harbored a combined 55 detected gonadal mosaic mutations— mostly single nucleotide mutations—with the potential to be transmitted to offspring (F01: 12, F02: 13, and F03: 30) (Figure 1b, Figure 1- figure supplement 1 and Supplementary Data). None were predicted to impair health when heterozygous, and they, thus, serve as likely neutral variants to model transmission. These mutations were present at AFs between 0.003 and 0.247 (mean: 0.047; standard deviation: 0.055), with those of lower AF typically restricted to sperm (‘*Sperm’*) and those of higher AF typically found in both sperm and blood or saliva (‘*Shared’*) (Figure 1c, and Figure 1- figure supplement 1). Their number and distribution were largely consistent with other unbiased analyses performed previously^10,13^. Of note, similar to our prior observations^13^, the sperm donor F01 (36 years of age) had an excessive number of soma-specific mutations (‘*Soma’*) at lower AFs, evidencing early clonal hematopoiesis^10,13^ (Figure 1- figure supplement 1).

For each sperm donor, eight, ten, or fourteen blastocysts, respectively (total: N=32), were assessed for transmission of paternal gonadal mosaic mutations. When calibrating our genotyping approach on whole-genome amplified blastocyst material, we found a false-negative rate of ∼0.06 and a false-positive rate of ∼0.0001 (Figure 2 – figure supplement 1). Across all 55 identified gonadal mosaic mutations, we observed 19 transmission events among 15 unique mutations (Figure 2 and Supplementary Data). For F02, two of the eight blastocysts did not show overall high quality in the MPAS for gonadal mosaic variants and thus were excluded from subsequent analyses.

**Figure 2.**
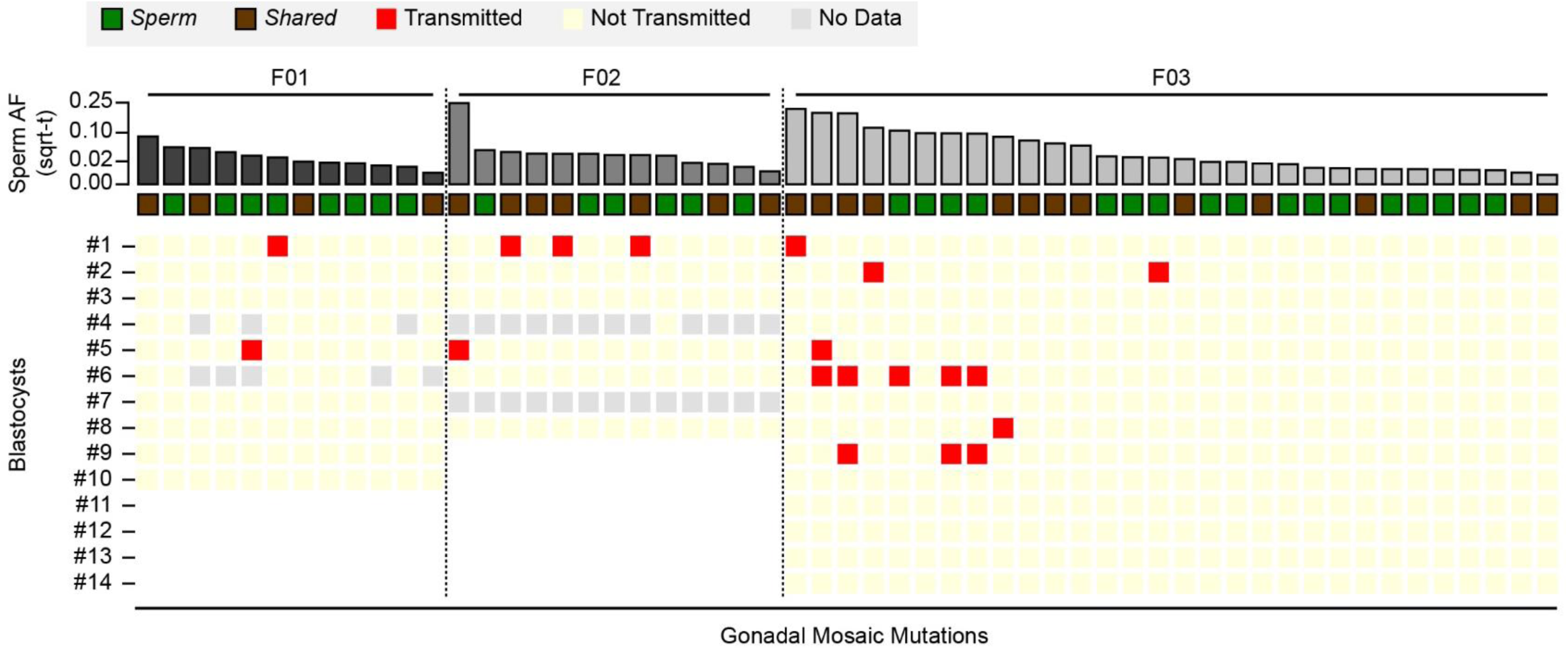
Transmission of gonadal mosaic mutations for each sperm donor. Mutations are ranked for each sperm donor by AF and visualized as square root-transformed (sqrt-t). Shown are the transmission of 12 mutations across 10 blastocysts (F01), of 13 mutations across 6 blastocysts (F02; 2 of 8 blastocysts did not show sufficient read depth across the mosaic mutations), and of 30 mutations across 14 blastocysts. In total, 19 transmissions of 15 unique mutations were observed. As expected, gonadal mosaic mutations of higher AF are more likely to transmit than those of lower AF. No Data: variant blastocyst pairs, for which read depth was below 20×.

Consistent with a model where the risk of transmission is proportional to sperm AF, mutations of higher AF evidenced higher blastocyst transmission rates than those of lower AF (R=0.67, p-value=2.9e-8, Pearson correlation, Methods). Although additional mutations may arise within earlier sperm lineages marked by prior mutations (e.g., blastocyst #2 in F03), somewhat unexpectedly, we observed co-segregation of mutations of almost equal AF (e.g., blastocyst #1 in F02 or blastocyst #6 in F03); this was reflected as a non-random distribution of transmitted mutations across blastocysts in F03 (Figure 2 – figure supplement 2). This suggests that some early male germ cell lineages or individual mitotic divisions are more susceptible to mutation than others, allowing stratification of sperm lineages into those with higher and those with lower mosaic load, and mutations from one sperm progenitor may uncouple during meiosis. Indeed, we observed different combinations of gonadal mosaic mutations across blastocysts (e.g., blastocysts #5, #6, and #9 in F03). This analysis demonstrates that *a priori* identified gonadal mosaic mutations have the potential to transmit to preimplantation blastocysts.

Based upon their individually measured AFs, we next calculated the expected number of transmission events across all blastocysts. Whereas for both F01 and F02 the observed transmission rate was within the 95% confidence interval of the expectation, for F03 and across all individuals when considered in total, the transmission was slightly below what was expected (Figure 3 and Figure 3 – figure supplement 1). This likely reflects a limitation of the model which assumes that gonadal mosaic mutations arise and transmit independently—a simplification, as mutations arising on the same lineage have the potential to co-transmit. This may be especially relevant if sperm lineages do not stochastically accumulate mutations during early development. Nevertheless, our observations closely reflect expectations, suggesting that gonadal mosaicism assessment can serve as a predictor of mutation transmission.

**Figure 3.**
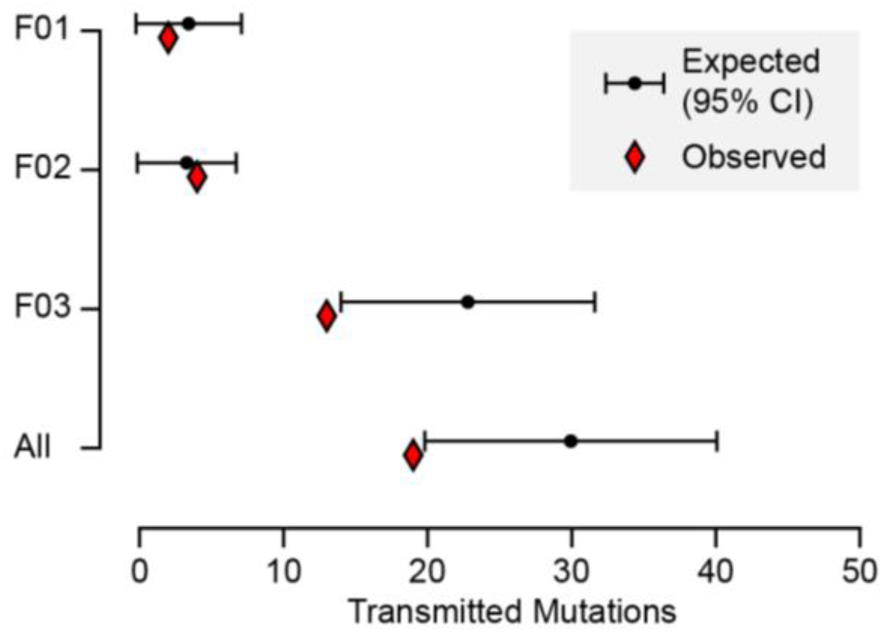
Expected and observed transmission of gonadal mosaic mutations to preimplantation blastocysts. The expected number of transmissions—based on the AF of all detected mutations in sperm and the number of analyzed blastocysts—is indicated with the mean and a 95% confidence interval for each sperm donor and across all sperm donors. Whereas F01 and F02 transmitted as expected, F03 showed a slight undertransmission, likely related to the non-independence of mutations due to shared lineage.

To validate the observation of predictable transmission and slight under-transmission, we further analyzed our previous gonadal mosaicism data from eight families with a total of 14 offspring, where we reported between 11-25 sperm-detectable mosaic variants per father^10,13^. We asked whether variants detected in sperm employing our unbiased detection pipeline transmitted to any of the 14 offspring. Across 131 sperm-detectable mosaic variants, we observed nine transmissions among seven unique variants (Figure 3 - figure supplement 2a and Supplementary Data). As this cohort had a lower number of observable transmission events per paternal sample due to the lower number of fertilization events (1-3 offspring compared to 8-14 blastocysts) and lower sequencing depth, we combined analysis of the eight families. The observed transmission rate was again slightly below the expected 95% confidence interval (Figure 3 - figure supplement 2b-c). This replication in live-born offspring supports the under-transmission observed in blastocysts and highlights the potential predictive power of gonadal mosaicism assessment.

Here we directly measured the abundance of gonadal mosaic variants and demonstrate transmission to preimplantation blastocysts for three couples undergoing IVF. These results provide a proof-of-concept that *a priori* detected gonadal mosaic mutations can transmit to blastocysts and therefore likely to offspring. If this approach is to be advanced in the clinic to prevent genetic disease, further research will be required: for instance, expanding the size of the cohort and directly assessing *a priori* identified pathogenic variants for transmission to blastocysts. In addition, further technological development will be required to enable the direct assessment of mutations from biopsies rather than whole blastocysts as implemented in this study.

We previously estimated that 1 in 15 males carry a potentially pathogenic mutation, detectable in approximately 5% of sperm cells^13^. While this likely only represents ∼15% of the monogenic sporadic component of diseases such as autism or congenital heart disease, this is likely the sole fraction that could be prevented with further advances. Thus, if these mosaic mutations were detected prior to pregnancy, and if mutation-carrying blastocysts were identified by PGT, there could be positive consequences for families through the prevention of pregnancy termination or pediatric diseases.

## Figures

**Figure 1 – figure supplement 1.**
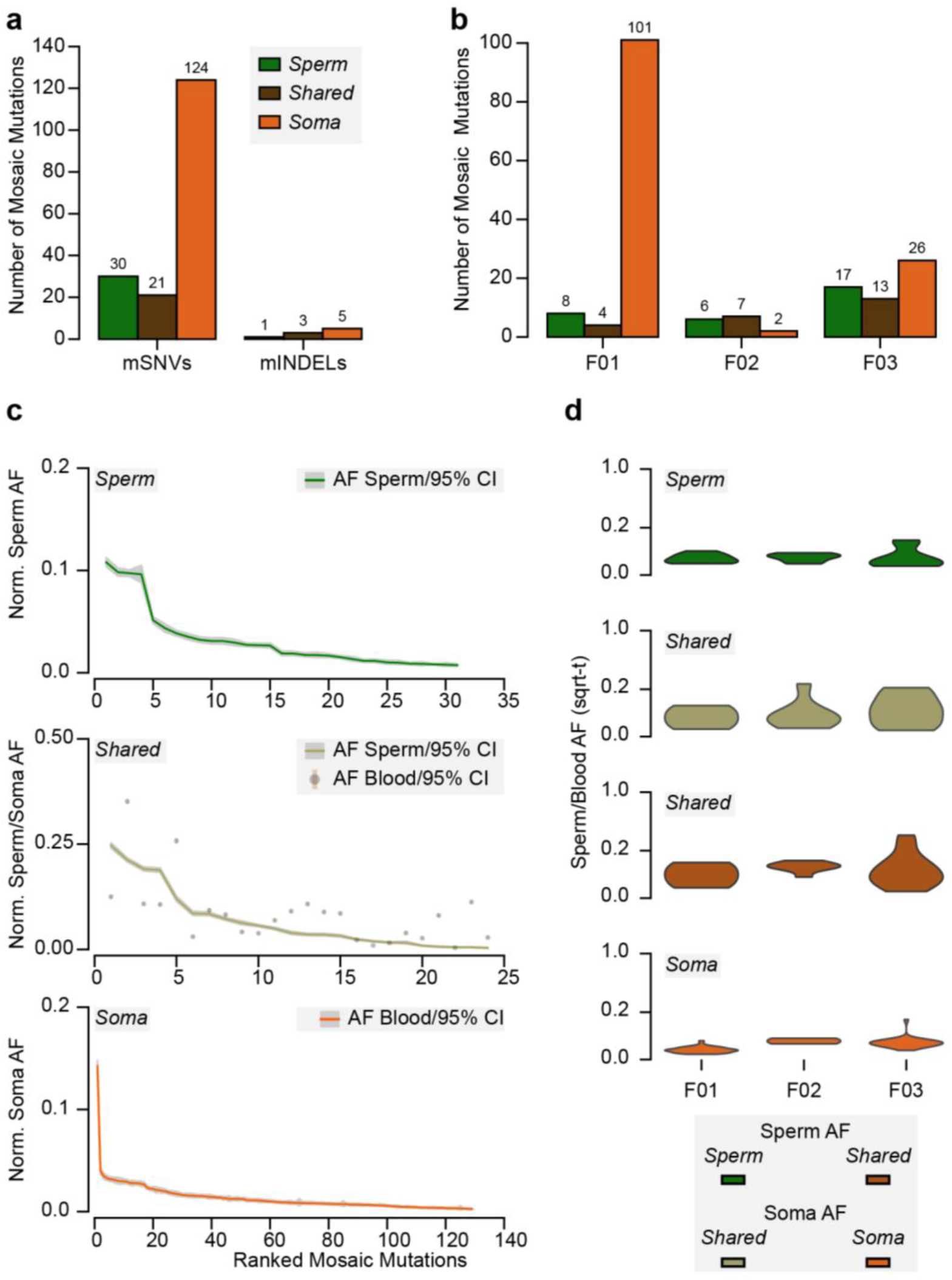
Detection of mosaic mutations in the three sperm donors. **(a)** Total number of identified mosaic single nucleotide mutations (mSNVs) and mosaic small insertions and deletions (mINDELs) across the three sperm donors. Mosaic mutations are divided into those detected only in sperm (‘*Sperm’*), those detected in both sperm and the somatic tissue (‘*Shared’*), and those detected only in the somatic tissue (‘*Soma’*; i.e., blood or saliva). **(b)** Number of mosaic mutations for each individual for each category. **(c)** Ranked plots for mosaic mutations across all sperm donors for each category. Shown are the allelic fractions (AF; normalized to chromosome counts) within the indicated tissue(s) and their 95% exact confidence interval. **(d)** Distribution of square root transformed (sqrt-t) AFs for each sperm donor for each category as a violin plot.

**Figure 2 – figure supplement 1.**
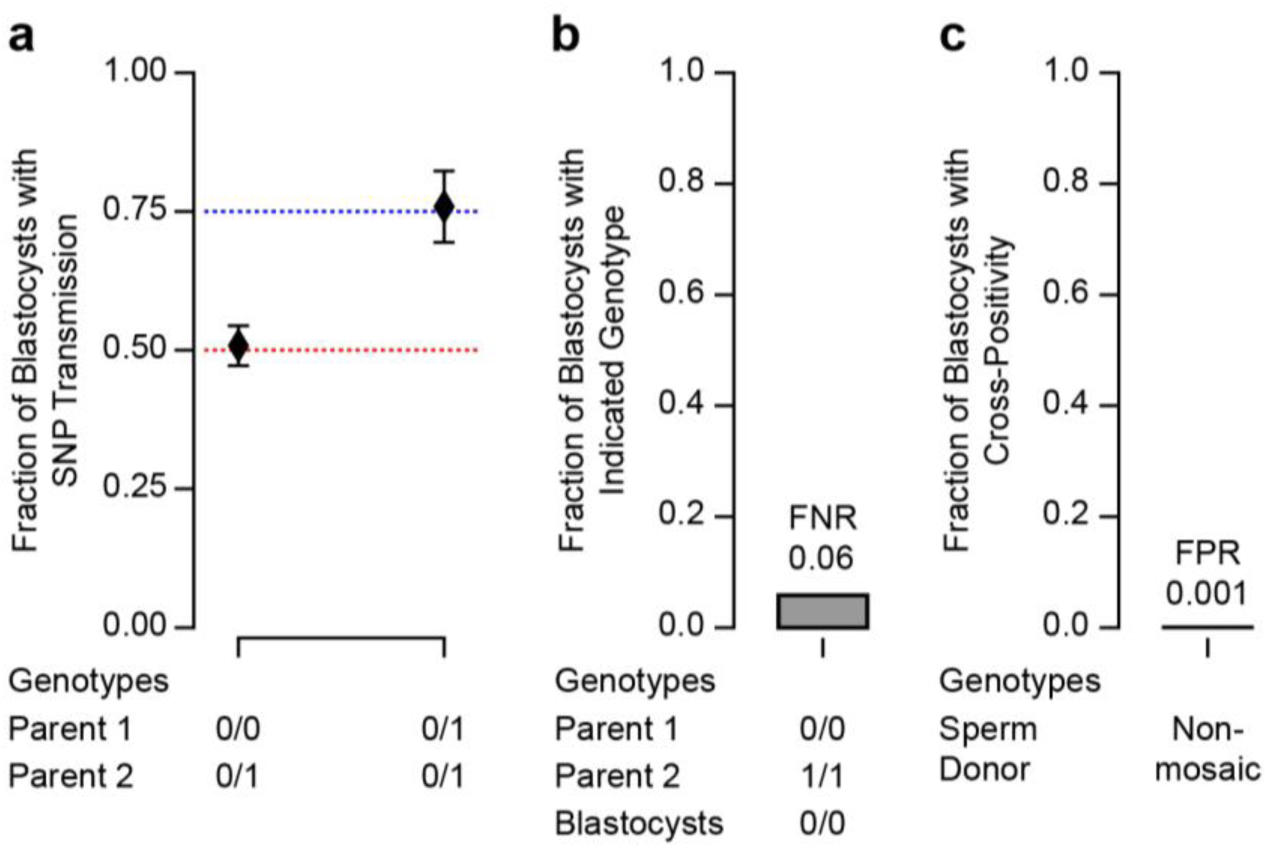
Determination of sensitivity and specificity for mutation detection in blastocysts. **(a)** Transmission of heterozygous single nucleotide polymorphisms (SNPs) detected in at least one parental genome. Parental genotypes are indicated as reference homozygous (0/0) and heterozygous (0/1). The 95% confidence interval of the observed fraction of transmissions overlaps with expectations of 0.5 (parents are heterozygous and reference homozygous, respectively) and 0.75 (parents are both heterozygous). 0/0 and 0/1: n=374 transmitted of 736 total; 0/1 and 0/1: n=129 transmitted of 170 total. As egg donor information was missing in F03, this analysis is restricted to F01 and F02. **(b)** Fraction of blastocysts that were determined to be negative when one of the parental genotypes is homozygous for an SNP (1/1). As a homozygous parent is expected to transmit the SNP to each blastocyst this allows us to determine the False Negative Rate (FNR=0.058). 0/0 and 1/1: n=81 transmitted of 86 total. As egg donor information was missing in F03, this analysis is restricted to F01 and F02. **(c)** Fraction of blastocysts that were determined to be positive when the mosaic mutation was identified in one of the unrelated sperm donors. These transmissions are unexpected and can only be explained by a technical artifact, allowing us to determine the False Positive Rate (FPR=0.001). Non-mosaic: n=1 ‘transmitted’ of 985 total.

**Figure 2 – figure supplement 2.**
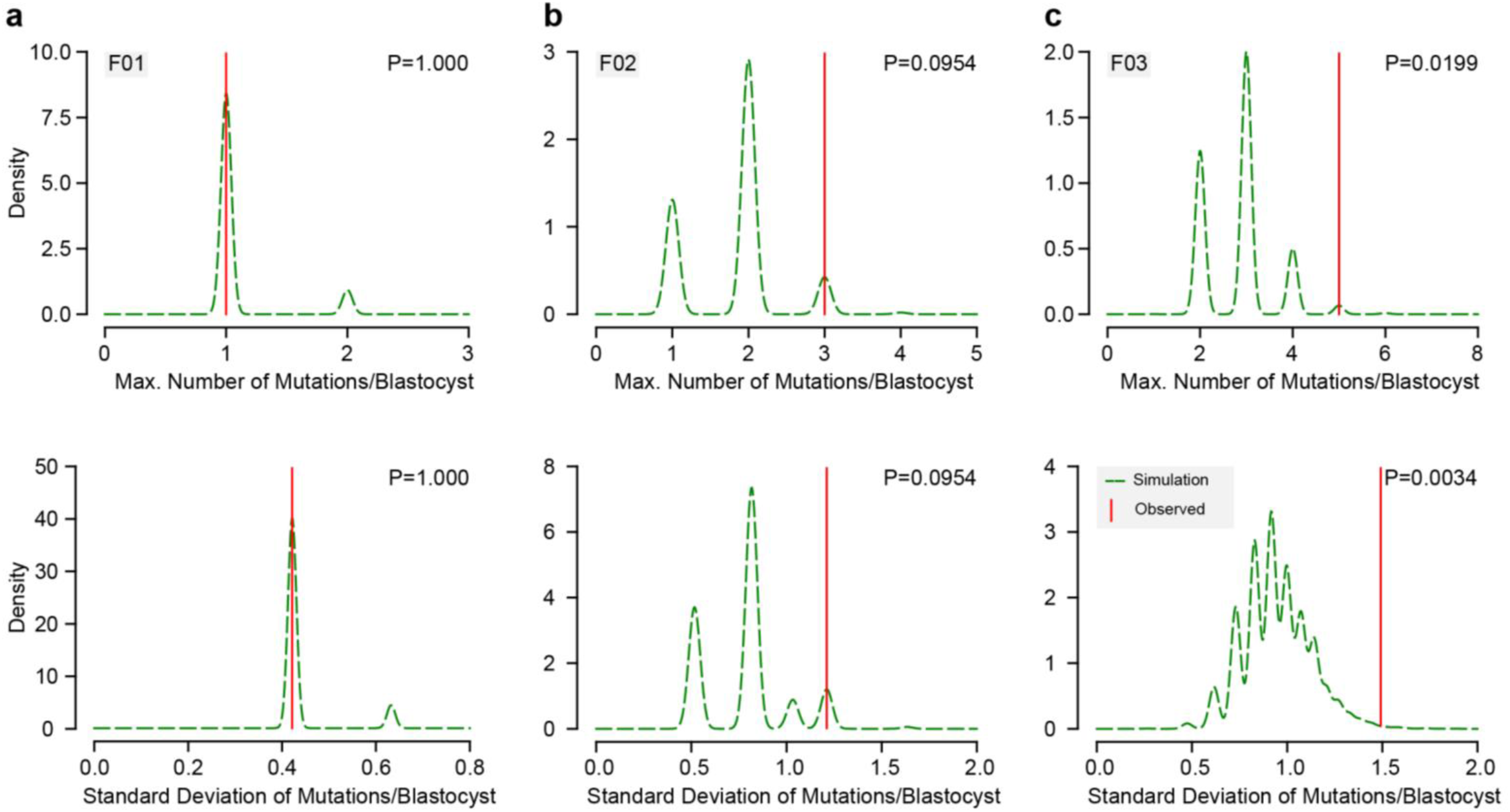
Evenness analysis of the distribution of transmitted mutations across blastocysts. **(a-c)** Permutation analysis for the observed number of transmitted mutations across blastocyst for F01 (a), F02 (b), and F03 (c), respectively. Underlying data points are either the maximum observed number of mutations within one blastocyst (top, Max. Number of Mutations/Blastocyst) or the standard deviation of the observed number of mutations across all Blastocysts (bottom, Standard Deviation of Mutations/Blastocyst) (n=10,000 permutations). Graphs show the density distribution of all permutations (green dashed line) and the corresponding observed value (vertical red line). Note that in F03, both the observed maximum number of transmitted mutations within one blastocyst and the standard deviation of observed transmissions across all blastocysts were significantly different from the random permutations (P<0.05).

**Figure 3 – figure supplement 1.**
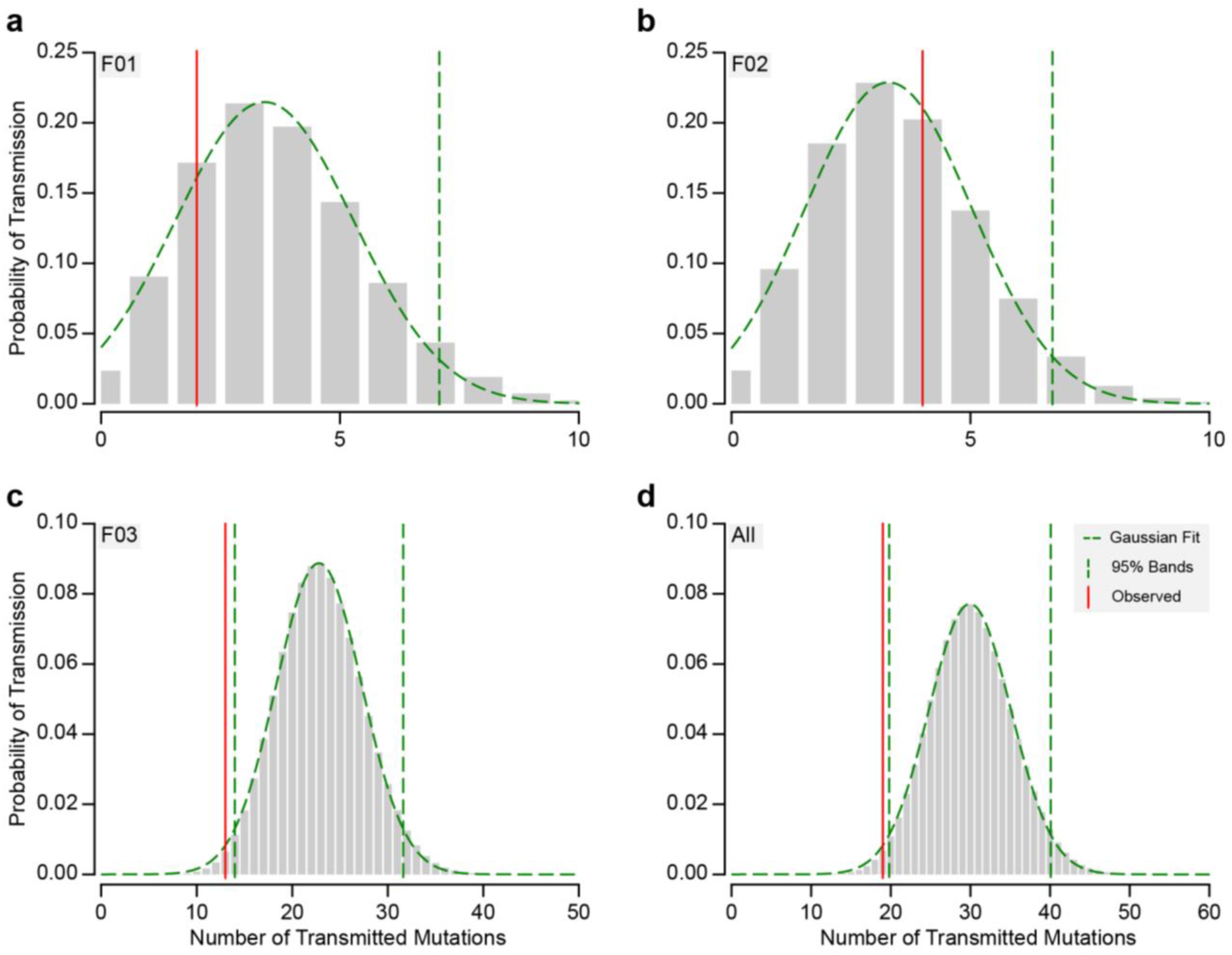
Detailed analysis of expected and observed transmission of gonadal mosaic mutations to preimplantation blastocysts. **(a-d)** Probability of the number of expected transmission events (grey bars), a Gaussian fit (green dashed line tracing the bars), the 95% confidence intervals if not 0 (vertical green dashed lines), and the observed number of transmission (vertical red line). This is shown for each sperm donor and blastocysts group (F01, F02, and F03 in a, b, and c, respectively) and across all (d). The mean and standard deviation determined from the Gaussian fit was used to generate data shown in Figure 3.

**Figure 3 – figure supplement 2.**
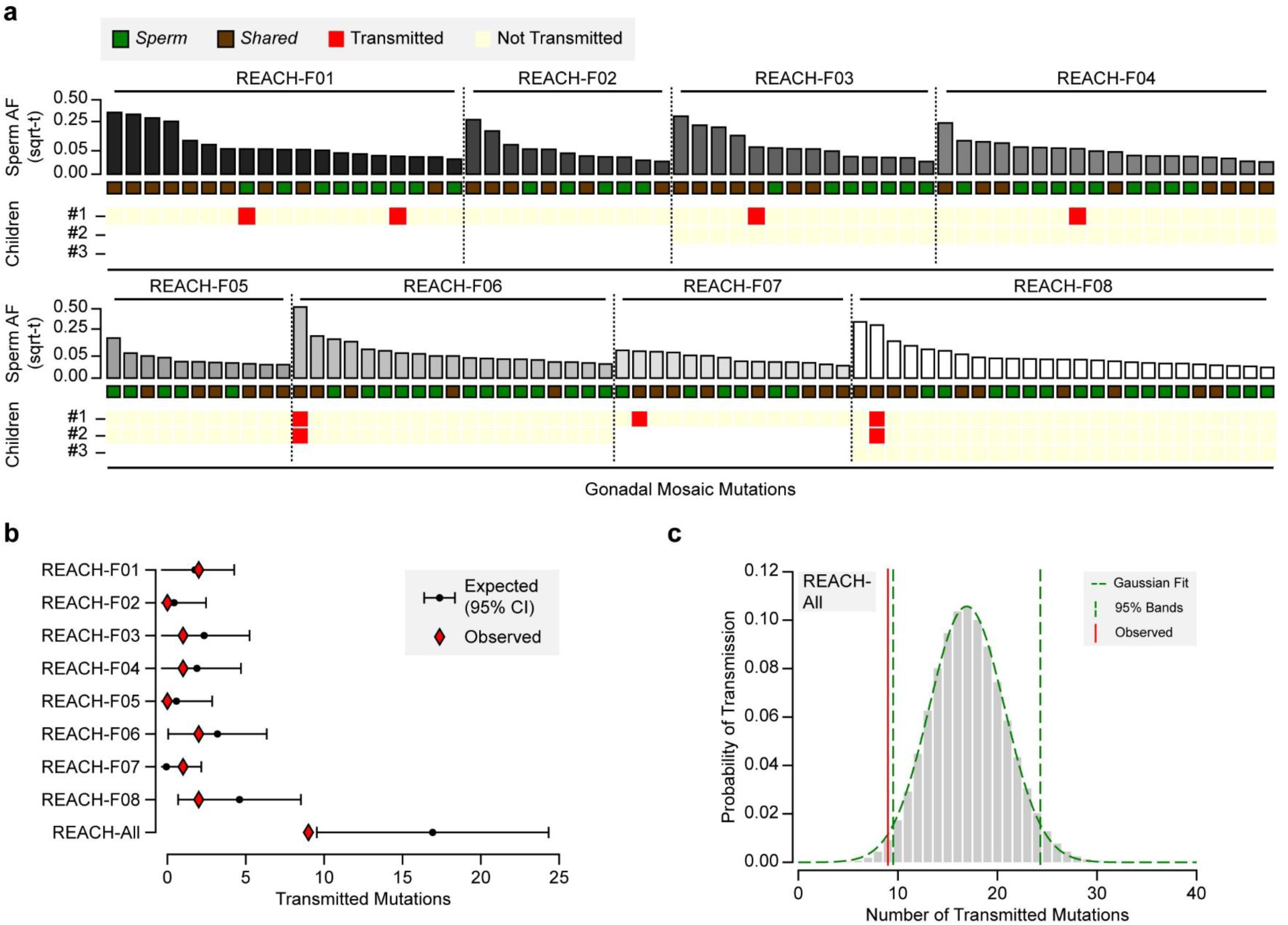
Transmission of gonadal mosaicism in eight previously described families. **(a)** Transmission of gonadal mosaic mutations for each of eight fathers whose gonadal mosaicism was previously described^10,13^. Each family had one to three offspring for whom variants were called from standard whole genome sequencing^16^. Mutations are ranked for each father by AF and visualized as square root-transformed (sqrt-t). Shown are the transmission of 19 mutations to 1 child (REACH-F01), of 11 mutations to 1 child (REACH-F02), 14 mutations across 2 children (REACH-F03), 18 mutations across 2 children (REACH-F04), 11 mutations across 2 children (REACH-F05), 19 mutations across 2 children (REACH-F06), 14 mutations to 1 child (REACH-F07), and of 25 mutations across 3 children (REACH-F08). In total, 9 transmissions of 7 unique mutations were observed. **(b)** Expected number of transmissions—based on the AF of all detected mutations in sperm and the number of analyzed children—is indicated with the mean and a 95% confidence interval for each father and across all fathers. Whereas, individually, each father transmitted within expectations, collectively, all fathers exhibited a slight undertransmission, comparable to what was found for blastocysts and F01-F03. **(c)** Probability of the number of expected transmission events across all fathers, visualized as in Figure 3 – figure supplement 1. The mean and standard deviation determined from the Gaussian fit was used to generate data shown in b.

**Figure 1 – table supplement 1.** Whole Genome Sequencing data depth.

| <u>Descriptor</u> |  | <u>Depth</u> |  |
| --- | --- | --- | --- |
| ID | Sample of origin | Mean | Standard Deviation |
| F01 | Sperm Donor's Sperm | 382.65 | 49.65 |
| F01 | Sperm Donor's Blood | 411.85 | 56.15 |
| F02 | Sperm Donor's Sperm | 295.00 | 46.00 |
| F02 | Sperm Donor's Saliva | 206.00 | 29.00 |
| F03 | Sperm Donor's Sperm | 308.18 | 47.18 |
| F03 | Sperm Donor's Saliva | 338.50 | 50.50 |

**Figure 1 – table supplement 2.** Massive Parallel Amplicon Sequencing data depth.

| <u>Descriptor</u> |  | <u>Depth</u> |  |
| --- | --- | --- | --- |
| ID | Sample of origin | Mean | Standard Deviation |
| F01 | Sperm Donor's Sperm | 19,313.71 | 6,174.02 |
| F01 | Sperm Donor's Blood | 22,070.96 | 6,816.18 |
| F01 | Egg Donor's Blood | 20,139.34 | 6,009.53 |
| F01 | Blastocyst #9 | 15,744.73 | 13,921.95 |
| F01 | Blastocyst #8 | 16,982.43 | 27,012.75 |
| F01 | Blastocyst #7 | 12,652.02 | 47,685.28 |
| F01 | Blastocyst #6 | 16,436.23 | 10,376.42 |
| F01 | Blastocyst #5 | 17,637.36 | 83,928.56 |
| F01 | Blastocyst #4 | 18,433.29 | 15,902.82 |
| F01 | Blastocyst #3 | 22,296.24 | 36,986.03 |
| F01 | Blastocyst #2 | 17,343.17 | 21,758.02 |
| F01 | Blastocyst #10 | 21,551.94 | 17,956.49 |
| F01 | Blastocyst #1 | 16,177.92 | 12,282.57 |
| F02 | Sperm Donor's Sperm | 13,572.89 | 4,078.05 |
| F02 | Sperm Donor's Saliva | 18,949.02 | 5,087.22 |
| F02 | Egg Donor's Saliva | 12,215.79 | 2,831.33 |
| F02 | Blastocyst #8 | 24,470.28 | 14,096.12 |
| F02 | Blastocyst #7 | 5.41 | 3.52 |
| F02 | Blastocyst #6 | 17,723.32 | 9,297.64 |
| F02 | Blastocyst #5 | 17,294.69 | 10,813.78 |
| F02 | Blastocyst #4 | 10.19 | 6.75 |
| F02 | Blastocyst #3 | 19,075.87 | 11,181.29 |
| F02 | Blastocyst #2 | 18,724.23 | 11,480.66 |
| F02 | Blastocyst #1 | 19,427.56 | 12,091.91 |
| F03 | Sperm Donor's Sperm | 17,296.13 | 8,575.26 |
| F03 | Sperm Donor's Saliva | 17,600.73 | 6,337.06 |
| F03 | Blastocyst #9 | 10,370.69 | 5,705.21 |
| F03 | Blastocyst #8 | 18,613.39 | 11,520.23 |
| F03 | Blastocyst #7 | 23,055.63 | 11,447.97 |
| F03 | Blastocyst #6 | 12,709.72 | 6,790.11 |
| F03 | Blastocyst #5 | 20,010.78 | 11,023.79 |
| F03 | Blastocyst #4 | 21,022.84 | 11,634.48 |
| F03 | Blastocyst #3 | 23,304.77 | 13,021.30 |
| F03 | Blastocyst #2 | 17,843.61 | 9,965.35 |
| F03 | Blastocyst #14 | 14,596.99 | 8,679.04 |
| F03 | Blastocyst #13 | 15,827.04 | 9,328.94 |
| F03 | Blastocyst #12 | 20,066.17 | 11,289.16 |
| F03 | Blastocyst #11 | 20,540.22 | 16,595.43 |
| F03 | Blastocyst #10 | 20,925.46 | 12,199.94 |
| F03 | Blastocyst #1 | 17,049.75 | 9,104.08 |
| Control | Control1's Blood | 17,588.46 | 5,261.91 |
| Control | Control2's Blood | 22,900.52 | 8,696.32 |

**Supplementary Data: Processed data tables.**

Provided as a separate file; legend contained within.

## Methods

### Donor Population, Recruitment, and Sample Preparation

For gonadal mosaicism assessment, we recruited three infertile couples (F01, F02, and F03) who had supernumerary blastocysts (minimum of 8) that wanted to donate them to research. The males (‘sperm donors’) all provided a fresh ejaculate and a somatic sample (blood for F01, saliva for F02, and F03). In accordance with reported population origin, the three fathers’ ancestries were most closely matched to Middle Eastern (F01) or European (F02 and F03) ancestry, employing nearest neighbor analysis of principle components^17^. One of the three females (‘egg donors’) was a non-identified egg donor (F01) and provided blood, another woman was the partner (F02) and provided saliva. The third chose not to provide a somatic sample (F03). DNA was extracted from each parental sample using documented procedures^10^, and extracted and amplified from vitrified blastocysts by thawing in phosphate-buffered saline supplemented with 5% bovine serum albumin and processing with REPLI-g whole genome amplification methods (Qiagen, Cat. #150343). Informed consent was obtained from all participants (custodians of the blastocysts) as well as from each participant in a study protocol approved by the University of California, San Diego IRB.

### Whole Genome Sequencing and Massive Parallel Sequencing

Whole genome sequencing (NovaSeq 6000, Illumina) of the sperm donor samples was performed to approximately 300× (Figure 1 - table supplement 1) as described^13^, then analyzed using the 300×MSMF (MuTect2, Strelka2, MosaicForecast) computational pipeline^13^. This pipeline demonstrates a 90% specificity and sensitivity of >95% for mutations at allelic fractions above 0.03 (Ref ^13^). All putative mosaic mutations (single nucleotide variants and small insertion-deletion mutations only) as well as 120 common and rare single nucleotide polymorphisms (SNPs), each detected as heterozygous in only one of the sperm donors, were then subjected to validation using massive parallel amplicon sequencing (MPAS, AmpliSeq Illumina Custom DNA Panel), to orthogonally assess each mutation in sperm donors, egg donors, and blastocysts. We also included two unrelated controls to control for false positive calls (Figure 1 – table supplement 2)^15^.

### MPAS Data Analysis

Mosaic mutations from tissues or blastocysts were confirmed or rejected based on the distribution of reference homozygous mutations and the signal in control samples at the same position as described previously^15^. The overall number and distribution of mosaic mutations were largely consistent with other unbiased analyses performed previously^10,13^. SNPs in non-blastocyst samples were assigned genotypes as reference homozygous, heterozygous, or alternate homozygous, based on the appearance of known genotypes previously determined in WGS data. Blastocysts were determined as genotype negative or positive for the SNPs.

To determine the expected number of transmissions to each blastocyst, the list of gonadal mosaic mutations and their measured AF in sperm for each sperm donor was used to determine the expected probability of transmission. Then, each blastocyst was combined with all other blastocysts from a single sperm donor or across all sperm donors as indicated. A gaussian 1D model was fitted using astropy’s LevMarLSQFitter and Gaussian1D to obtain the mean as well as the standard deviation and 95% confidence interval. Code used for data analysis and generation of all plots can be found on GitHub: https://github.com/shishenyxx/Sperm_transmission_mosaicism. Blastocysts with less than 10% detectable genomic positions according to the panel were excluded from the downstream analysis.

### Determination of False Negative and False Positive Rate of Transmission to Blastocysts

To calibrate our genotyping approach on whole-genome amplified blastocyst material, we determined the transmission of heterozygous variants from sperm and egg donors (Figure 2 – figure supplement 1). We found that transmission followed expected genetic patterns; furthermore, homozygous variants from a parent—which should be present across all blastocysts—were transmitted in approximately 94% of those analyzed, suggesting a false negative rate of ∼0.06. Importantly, this false-negative rate also includes potential allelic dropout, which can be problematic for single-cell studies or amplification from biopsies.

Conversely, when assessing the presence of gonadal mosaic mutations identified in one sperm donor in the blastocysts of the other two, we only found one such event out of 985 possible, suggesting an MPAS false positive rate of ∼0.001. Thus, this approach provides sensitivity to detect clinically relevant gonadal mosaicism (i.e., mutations with a measurable and potentially actionably abundance) and specificity to assess transmissions of perm mosaic mutations to blastocysts, and it suggests that whole genome-amplified blastocysts exhibit modest allelic dropout, even though allelic imbalance was frequently observed.

### Determination of Mosaicism in MPAS data

For each mosaic mutation or SNP, we determined the estimated allelic fraction (AF) and its 95% confidence interval based on MPAS. As a baseline for observed noise, we determined the distribution of reference homozygous SNPs and their lower 95% confidence interval (population threshold). Mutations considered as a mosaic in a sperm donor’s tissue fulfilled three criteria: 1] read depth for the position was at least 100x; 2] the observed 95% confidence interval of the mutation did not overlap with the population threshold or the upper 95% confidence interval of the control samples; 3] either control’s lower 95% confidence interval limit had to be below the population threshold. For a blastocyst, to be considered positive for the mutation, similar criteria were applied, but the lower 95% confidence interval could not overlap with 0.05 AF and the read depth at this position had to be equal or above 20x. For SNPs in blood, saliva, and sperm, a mutation was considered heterozygous if above or equal to 0.2 and below 0.8 AF; it was considered homozygous if above or equal to 0.8 AF. Blastocysts were considered positive (either heterozygous or homozygous, not distinguished) if above 0.05 AF.

### Determination of Evenness of Transmitted Mutations across Blastocysts

For each of F01, F02, and F03, the observed number of transmitted mutations were randomly assigned to the different blastocysts in 10,000 permutations. For each permutation, the maximum number of mutations transmitted to one blastocyst as well as the standard deviation of the number of mutations transmitted across each blastocyst were determined. The obtained distribution across the 10,000 permutations was then compared to the observed value. A permutation P-value was calculated based on the tail probability (the number of permuted values larger than or equal to the observed value over the total number of permutations).

### Distribution and Impact of Gonadal Mosaic Mutations

The 55 gonadal mosaic mutations were distributed across all chromosomes—except for chromosomes 19, 22, and X/Y—roughly as expected based on chromosomal length. As expected based on prior work, the vast majority was found in intergenic (n=28) or intronic (n=24) regions; one mutation each was found in a 5’-UTR, the intron of a non-coding RNA, and an exon. The exonic mutation (F03, AF=0.063) resulted in a non-synonymous SNV in RABGAP1 (NM_012197; p.Asp74Gly), a rare, previously reported mutation (allele frequency of 7.96e-6) with no known disease association.

### Reanalysis of the REACH cohort

Re-analysis of the REACH cohort data was performed by combining the list of detected gonadal mosaic mutation^*10,13*^ with previously established variants called from the trio WGS^*16,18*^. For each gonadal mosaic mutation, we determined the occurrence in a child similar to what was done for blastocysts. Finally, the expected probability of transmission was determined as described above.

## Data availability

Raw whole-genome sequencing data of the bulk sperm, saliva, and blood are available on SRA under accession numbers PRJNA753973 and PRJNA588332. The raw genotyping table is provided as Supplementary Data 1. The code for data analysis and statistical tests is available on Github (https://github.com/shishenyxx/Sperm_transmission_mosaicism).

## Acknowledgments

We thank Drs. Yan Ding, Shareef Nahas, and Lucitia van der Kraan (Rady Children’s Institute for Genomic Medicine, San Diego) for sequencing and computational support, and Drs. Louise Laurent and Jonathan Sebat (University of California, San Diego) for feedback on the manuscript.

## Author contributions

M.W.B., X.Y., and J.G.G designed the study. M.W.B., X.Y., J.G.G., and A.J.M. designed the experiments. M.W.B., X.Y., and J.M.-V. performed experiments. X.Y., X.X., and M.W.B. analyzed the data. A.J.M., V.S., and J. M.-V. contributed to participant recruitment and clinical data acquisition. M.W.B. and J.G.G. supervised all parts of the study. M.W.B., X.Y., and J.G.G. wrote the manuscript. All authors approved the final version of the manuscript.

